# Reconstruction of metagenome-assembled genomes from aquaria

**DOI:** 10.1101/2021.05.28.446213

**Authors:** Cassandra L. Ettinger, Jordan Bryan, Sima Tokajian, Guillaume Jospin, David Coil, Jonathan A. Eisen

**Affiliations:** Genome Center, University of California, Davis, CA, USA; Department of Evolution and Ecology, University of California, Davis, CA, USA; Department of Microbiology & Plant Pathology, University of California, Riverside, CA, USA; College of Agriculture and Life Sciences, Cornell University, Ithaca, NY, USA; Department of Natural Sciences, Lebanese American University, Byblos, Lebanon; AnimalBiome, Oakland, CA, USA; Department of Medical Microbiology and Immunology, University of California, Davis, CA, USA

**Keywords:** aquarium, metagenomics, metagenome assembled genomes, candidate phyla radiation

## Abstract

We report 11 metagenome-assembled genomes (MAGs) reconstructed from freshwater and saltwater aquaria including representatives of Polynucleobacter, Anaerolinae, Roseobacter, Flavobacteriia, Octadecabacter, Mycobacterium and Candidate Phyla Radiation (CPR) members. These MAGs can serve as a resource for aquatic research and elucidating the role of CPR taxa in the built environment.

## Announcement

Microbial communities play critical roles in aquaria health. Aquaria support complex multi-trophic interactions between fish, invertebrates, plants and microbial communities that occur in an enclosed built environment. Understanding the genomics of aquaria microbial communities is critical to understanding the health of other enclosed aquatic systems.

Samples were collected prior to the start of an undergraduate research project that investigated microbial community assembly of multiple aquaria in the Fall of 2012 at the University of California, Davis (1). Tropical tank sediment (n=3), cold reef tank sediment (n=1), freshwater tank sediment (n=3), cold reef tank water (n=3), freshwater wipes (n=3) and freshwater tank water (n=3) were collected and processed for DNA extraction as described in Bik et al (1). Libraries were made using a Nextera XT DNA library sample preparation kit (Illumina, Inc.) and were sequenced on an Illumina MiSeq (paired end, 150 bp reads).

All reads were co-assembled using MEGAHIT (2) v.1.0.6. Metagenome-assembled genomes (MAGs) were generated using anvi’o v. 2.3.2 (3). First, a contig database was produced using ‘anvi-gen-contigs-database’ and open reading frames identified with Prodigal (4) v.2.6.2. We then used ‘anvi-run-hmms’ to run HMMER v3.1b2 (5) to identify bacterial (6) and archeal (7) single-copy genes. Contig taxonomy was inferred using Kaiju v.1.5.0 (8) with the NCBI BLAST non-redundant protein database including fungi and microbial eukaryotes v.2016-09-18. Reads were mapped using Bowtie2 v.2.2.8 (9) and samtools v.1.4.1 (10). Using ‘anvi-profile’ and ‘anvi-merge’, contigs > 2.5 kbp were mapped to samples and then profiles were combined. On average, 780,565 reads per sample mapped to the contig database with the majority of mapped reads from cold reef tank water (57.3%) and freshwater tank water (41.7%). Contigs were clustered using ‘anvi-cluster-with-concoct’ to automatically bin MAGs (11). MAG completeness and contamination was assessed in anvi’o using ‘anvi-summarize’ and confirmed with CheckM v.1.0.7 (12). Phylosift v. 1.0.1 (13) was used to place MAGs in a phylogenetic context to provide additional information about taxonomic assignments. Candidate Phyla Radiation (CPR) genomes were identified with ‘anvi-script-gen-cpr-classifier’ and ‘anvi-script-predict-cpr-genomes’ using the Brown et al (14) and Cambell et al (6) databases. CPR genome completion was then estimated for 43 single copy marker genes (14). The CPR are putatively a diverse group of uncultured bacterial lineages with poorly understood metabolic functions known mostly from metagenomic sequencing work. Representatives of CPR have been previously found in a wide range of aquatic habitats including bioreactors, ocean, lakes, groundwater and waterways (15– 21).

We report two high-quality draft MAGs >90% completion and four medium-quality draft MAGs with >70% completion (Table 1). Additionally, we report five draft MAGs that were identified as potential CPR genomes with >90% completion (Table 2). These metagenome-assembled genomes will enable deeper insights into the ecology of aquaria microbial communities and also into the possible functional roles of understudied lineages (e.g. CPR members) in the built environment.

**Table 1.**
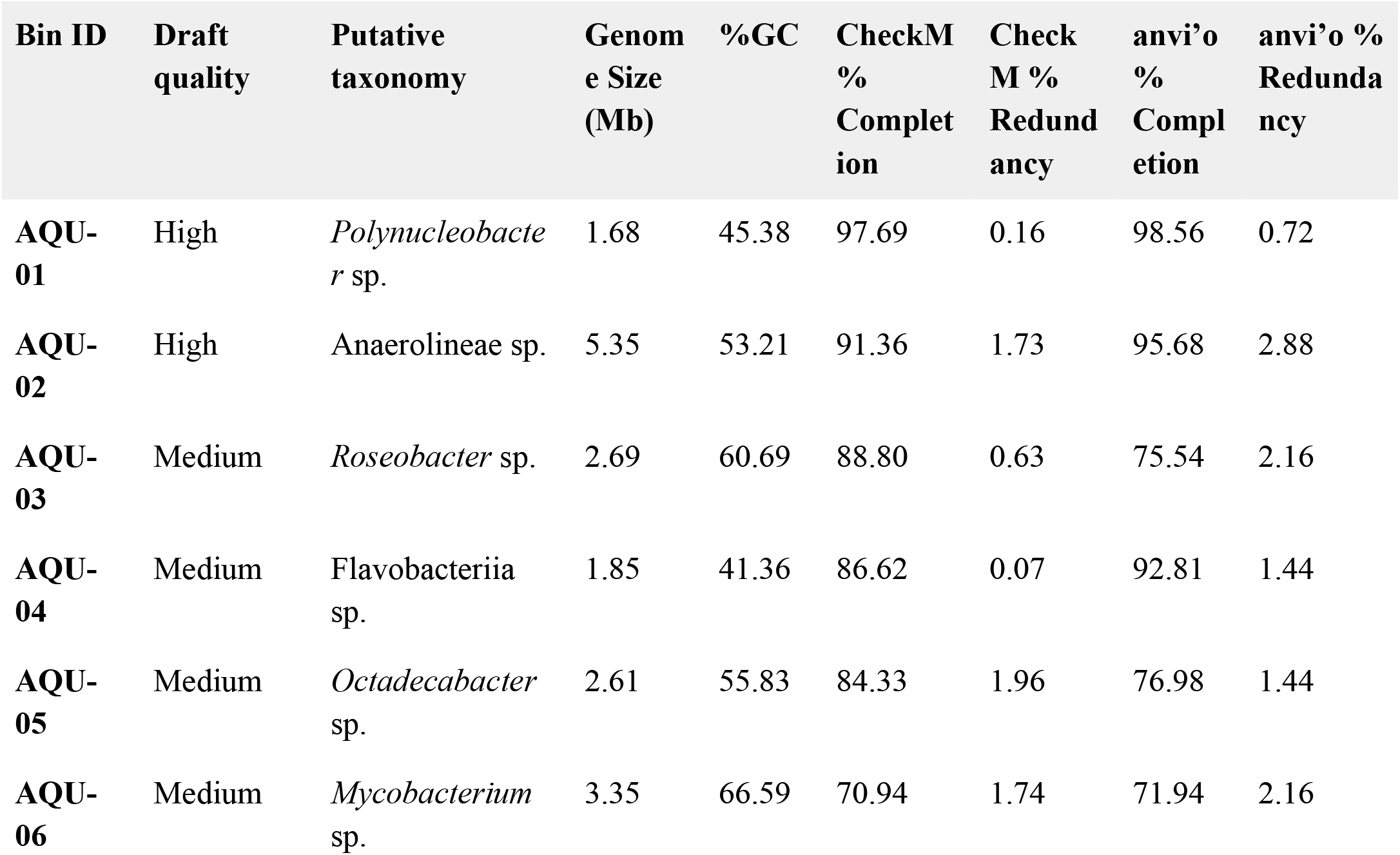
Genomic feature summary for metagenome-assembled genomes identified from aquaria.

| Bin ID | Draft quality | Putative taxonomy | Genome Size (Mb) | %GC | CheckM % Completion | CheckM % Redundancy | anvi'o % Completion | anvi'o % Redundancy |
| --- | --- | --- | --- | --- | --- | --- | --- | --- |
| AQU-01 | High | *Polynucleobacter* sp. | 1.68 | 45.38 | 97.69 | 0.16 | 98.56 | 0.72 |
| AQU-02 | High | Anaerolineae sp. | 5.35 | 53.21 | 91.36 | 1.73 | 95.68 | 2.88 |
| AQU-03 | Medium | *Roseobacter* sp. | 2.69 | 60.69 | 88.80 | 0.63 | 75.54 | 2.16 |
| AQU-04 | Medium | Flavobacteriia sp. | 1.85 | 41.36 | 86.62 | 0.07 | 92.81 | 1.44 |
| AQU-05 | Medium | *Octadecabacter* sp. | 2.61 | 55.83 | 84.33 | 1.96 | 76.98 | 1.44 |
| AQU-06 | Medium | *Mycobacterium* sp. | 3.35 | 66.59 | 70.94 | 1.74 | 71.94 | 2.16 |

**Table 2.** Genomic feature summary for Candidate Phyla Radiation (CPR) metagenome-assembled genomes identified from aquaria.

| Bin ID | Putative taxonomy | Genome Size (Mb) | %GC | CPR % Completion<sup>a</sup> |
| --- | --- | --- | --- | --- |
| AQU-07 | Candidatus Shapirobacteria sp. | 0.86 | 35.61 | 93.02 |
| AQU-08 | Candidatus Kerfeldbacteria sp. | 1.07 | 46.20 | 93.02 |
| AQU-09 | Candidatus Uhrbacteria sp. | 1.12 | 51.31 | 93.02 |
| AQU-10 | Candidatus Moranbacteria sp. | 0.89 | 43.56 | 93.02 |
| AQU-11 | Candidatus Saccharibacteria sp. | 1.09 | 47.69 | 90.70 |

<sup>a</sup>Completion estimates were generated using 43 single copy markers for CPR following Brown et al (14).

## Data availability

The raw sequencing reads, co-assembly and individual MAGs were deposited at DDBJ/ENA/GenBank under BioProject accession number <u>PRJNA728121</u>. Contigs identified as possible contaminants or adaptors by NCBI’s Contamination Screen were subsequently trimmed or removed from the co-assembly or individual MAGs during deposition.

## Acknowledgements

Illumina sequencing was performed at the DNA Technologies Core facility in the Genome Center at the University of California, Davis, Davis, California (UCD). We acknowledge the contributions of multiple people for help in various stages of the project, especially Thein Mai and Matt Wein for access to and assistance with the aquaria used in this study. Funding for this study was provided by the Alfred P. Sloan Foundation through their program in the “Microbiology of the Built Environment”.

## Notes

### Competing Interest Statement

The authors have declared no competing interest.

